# Salivary miRNA and microbial profiles reflect different responses to psychosocial stress

**DOI:** 10.64898/2026.03.20.713173

**Authors:** Sergio Garbarino, Nicola Magnavita, Barbara Pardini, Sonia Tarallo, Fabrizio Cipriani, Alessandro Camandona, Giulio Ferrero, Egeria Scoditti, Alessio Naccarati

**Affiliations:** Department of Neurosciences, Rehabilitation, Ophthalmology, Genetics and Maternal/Child Sciences (DINOGMI), University of Genoa, Largo Paolo Daneo 3, 16132 Genova, Italy; Occupational Epidemiology and Health Unit, Department of Life Sciences and Public Health, Università Cattolica del Sacro Cuore, Rome, Italy; Genetic and Molecular Epidemiology Unit, Italian Institute for Genomic Medicine (IIGM), c/o INOC – Istituto Nazionale Oncologico Candiolo, IRCCS, viale della ricerca 7, 10060 Candiolo, Italy; Department of Clinical and Biological Sciences, University of Turin, Regione Gonzole 10, 10043, Orbassano, Italy; CNR Institute of Clinical Physiology, Via Giuseppe Moruzzi, 1, 56124, Pisa, Italy

## Abstract

**Background:** Psychosocial stress is a risk factor for mental and physical illness. Previous studies have shown the potential of microRNAs (miRNAs) and the microbiome as biomarkers and/or mediators for stress responses. Here, we aimed to identify changes in these molecular markers in saliva samples from a cohort of 113 male police officers in service, stratified by stress response (SR) level (low, intermediate, and high), as a reference model for psychosocial stress.

**Methods:** Salivary miRNome profiles and microbiome composition were analyzed by small RNA sequencing and shotgun metagenomics, respectively. SR-related miRNAs were identified with Docker4Seq and DESeq2 and functionally characterized using RBiomirGS. Differences in microbial communities among study groups were analyzed using MetaPhlAn, and microbial pathway prediction was performed with HUMAnN. A multi-omics integration of miRNA and microbial profiles was performed with the mixOmics framework.

**Results:** Eighteen miRNAs were dysregulated (adj. p<0.05) between high- and low-SR groups and showed a progressive alteration from low- to high-SR groups. Functional enrichment analysis indicated that dysregulated miRNA targets were involved in apoptosis, cellular stress responses, and metabolic regulation. Despite limited shifts in oral microbial composition and diversity among the SR groups, the high-SR group showed a significant (adj. p<0.1) increase in microbial species involved in inositol degradation pathways, as well as a decrease in bacteria involved in L-tryptophan and thiamine biosynthesis, whose altered levels distinguished the high-SR group.

**Conclusions:** Salivary miRNAs and oral microbiota may serve as non-invasive indicators of SR, helping to understand chronic psychosocial stress and cellular and behavioral responses.

**Plain language summary:** Psychosocial stress is a major risk factor for mental and physical illness, with emerging evidence pointing to oral miRNAs and the microbiome as potential biomarkers. This study investigated stress-associated molecular changes in saliva from 113 male police officers stratified by perceived stress response (SR) into low, intermediate, or high responders. Eighteen miRNAs were dysregulated between high- and low-SR groups, whose target genes were involved in apoptosis, cellular stress responses, and metabolic regulation. Despite limited shifts in microbial populations, a functional analysis indicated increased taxa involved in inositol degradation and reduced taxa involved in L-tryptophan and thiamine biosynthesis, distinguishing high-SR individuals. These findings suggest that salivary miRNAs and microbiota may serve as non-invasive indicators of psychosocial stress, whose altered levels could reflect different SR.

## Introduction

Stress is an engineering term that refers to the tension or load to which a material is subjected. In medicine and biology, it represents the stimulus that elicits the general homeostatic response in living organisms^1,2^. This response to real or perceived environmental, psychological, or physiological threats triggers coordinated neuro-immuno-endocrine processes to maintain or restore homeostasis^2^. Depending on stress intensity, duration, predictability, and individual coping capacity, the stress response (SR) may elicit distinct physiological and behavioral reactions. These responses could be adaptive, leading to habituation (eustress), or ineffective and even maladaptive, such that the organism fails to return to physiological and/or psychological homeostasis (distress) and may be exposed to altered mental and physical health^3^. It is well known that there can be stress without distress^4^: in other words, exposure to the same stressor can have divergent effects in different individuals, being positive for one, neutral for another, and negative for a third. Modern theories of stress focus on people’s interactions with their work environment, on the psychological mechanisms underlying these relationships, or on the transactional mechanisms of cognitive appraisal and coping. Distress is not yet a disease, but it can become one. Indeed, when SR demands exceed available resources, there could be an increase in the risk for the development of acute or chronic diseases, including infection, cardiovascular, metabolic, and mental diseases, as well as cancer^5^.

Advances in understanding the basic regulatory mechanisms of SR have revealed the role of microRNAs (miRNAs), a class of small non-coding RNAs with a pivotal role in the post-transcriptional regulation of gene expression^6,7^. miRNAs can be detected in extracellular fluids, including serum, plasma, and saliva. Accumulating evidence indicates aberrant miRNA expression in several human diseases^6^; therefore, miRNAs may serve as potential biomarkers and signaling molecules in pathogenic pathways. The role of miRNAs in stress signaling was recognized early in loss-of-function studies, in which experimental animals with miRNA knockout or inactivation developed normally but were unable to cope with stress^8^. Human studies have confirmed the involvement of miRNAs in the regulation of acute and chronic SR^6,7,9–12^, underscoring their potential to improve understanding and monitoring of stressful conditions.

At the same time, thanks to deep sequencing and sophisticated computational analyses, the ability to comprehensively investigate the human microbiome has, over the last decade, enabled numerous studies on the relationship between microbial species and disease^13^. Research has also revealed many associations between gastrointestinal tract microbes and stress/SR^14^. The gut-brain axis hypothesis has been corroborated by direct interactions between intestinal microbes and changes in mental health status, specific disorders, or general stress. More recently, the study of the oral microbial composition has emerged as important for investigating the relationships among multifaceted aspects of stress, the host, and the microbes that interact with it^15,16^.

Work-related distress is a chronic psychosocial condition that occurs “when the demands of the work environment exceed the employees’ ability to cope with (or control) them”^17^, or when there is a discrepancy between the efforts made to work and the rewards received for the work done^18^. Distress is the second most common work-related health issue in Europe, leading to anxiety, depression, fatigue, worse quality of life, physical illness, as well as reduced participation, performance, and safety at the workplace^19^.

Peculiar groups of workers, such as public safety personnel and first responders, including police, firefighters, and emergency dispatchers, are at high risk of potential work-related mental health issues as a consequence of psychosocial distress^20^. Police officers may be significantly exposed to both operational work-related challenges (e.g., violence and trauma, work shift, working hours) as well as to organizational stressors (e.g., negative public perception of police, perceived lack of departmental support, interface with the judicial system, bureaucracy)^21^. Greater perceived work stress was associated with a higher risk of developing or aggravating mental and physical health problems, including sleep problems, anxiety, depression, musculoskeletal disorders, metabolic syndrome, cardiovascular disease and cancer^22–25^. An umbrella review, covering data on police officers from 43 countries, identified twenty-six adverse health outcomes and 220 underlying risk factors, 136 of which were modifiable, including work-related stress^26^. Moreover, work-related stress also reduced well-being^27^ and work performance, thus threatening public safety^28^.

The present study aimed to evaluate the association between chronic psychosocial SR, salivary miRNA profiles, and the oral microbiome. As a model of psychosocial stress, we leveraged the longitudinal Genoa Police Cohort, a census of workers in service in the mobile department of Genoa (Italy) in 2009, which was monitored for SR over thirteen years until 2022^29^. Our hypothesis was that individuals grouped by SR (low, intermediate, and high distress) also differ in their salivary miRNome and microbiome, reflecting their different levels of resilience.

## Methods

### Study cohort

The present study was a cross-sectional analysis conducted in 2022 on the Genoa Police Cohort composed of police workers from the “VI Reparto Mobile” (Mobile Unit) of Genoa (Italy), who were in service in 2009 (n=292) and remained in the same unit until 2022 (n=113). Members of the mobile departments are specialized and responsible for maintaining public order and performing work widely recognized as physically and psychologically demanding. They carry out the same public-order tasks regardless of their qualifications, rank, or seniority. Workers must maintain strict control of their lifestyle, and their preparation includes a program of physical exercises and training for the use of firearms. Service hours may vary depending on service needs, and it is common for members of these units to work a high number of overtime hours, even at night. Exposure to acts of violence is inherent to professional commitment. The health monitoring and promotion project for the police officers was authorized by the Italian Ministry of the Interior in agreement with all the workers’ trade unions.

The control of the workers’ mental health status (e.g., stress) was delegated to a university professor (N.M.) external to the administration, to ensure that any findings of stressful conditions would not have a negative impact on the careers or earning capacity of the workers. Workers participated in the different phases of the program at high rates. The study protocol was approved by the Ethics Committee of the Università Cattolica del Sacro Cuore of Rome, Italy (approval n. 285, 16 July 2020), and conducted in accordance with the Declaration of Helsinki. All participants provided their written informed consent to participate in the study. Since only two female workers were recruited, they were excluded from the analyses reported in this study for statistical reasons. Current smokers as well as subjects with periodontal problems were excluded. Sociodemographic and work-related characteristics, including age, education, marital status, presence of offspring, type of housing, military rank, work experience, and past medical history, were collected through interviews.

### Stress assessment

Accurate measurement of chronic occupational stress requires addressing several challenges. Stress measured in an occupational cohort cannot be compared with that of the general population, which is not exposed to occupational stressors. Consequently, occupational studies require internal control. The subjective nature of the SR makes it difficult to establish the cut-off score of questionnaires. Furthermore, a worker’s perception of occupational stress is inevitably influenced by life events and may therefore vary over time in response to such unpredictable events. To minimize these problems, workers from the Genoa cohort were compared with themselves, examining the distribution of police officers’ responses to a homogeneous occupational stressor, namely public order tasks.

The questionnaires were administered multiple times each year of observation, so that any external event could affect only the measurement closest in time, with little effect on the overall classification. Work-related stress was measured using two models to capture the different aspects of interactions between professional conditions and individuals that arise in relation to different operational situations^30^. The Italian versions^31^ of the short forms of the Karasek Demand/Control questionnaire^32^ and the Effort/Reward Imbalance questionnaire^18^ were administered. The distribution of scores for each questionnaire administered during the first decade of observation was divided into quartiles, and each worker was assigned a score from 1 to 4 corresponding to their quartile. The sum of the scores yields an ordinal measure, “Stress Response” (SR), with values ranging from 1 to 40. Higher SR values indicate a greater response to homogeneous stressors in public order control^29^.

Leaving the Mobile unit cohort can occur for supervening age limits and retirement, but more frequently, the dropout can occur due to the reporting of health problems that lead to moving to other forms of state service. During the observation period, the percentage of workers in the highest SR who continued to serve in the unit decreased more than that of the most resilient workers. Moreover, the subdivision of workers according to their resilience level allowed us to verify that those belonging to the extreme groups differed in terms of personality traits^33^, levels of anxiety and depression^22^, absenteeism rates^34^, quantity and quality of sleep^35^, and metabolic syndrome incidence^23^.

### Saliva collection and nucleic acid extraction

Saliva samples were collected from January to March 2022 from 113 subjects using the Isohelix GeneFiX™ DNA/RNA Saliva collectors kit (Isohelix), following the manufacturer’s instructions. Participants were asked to refrain from eating for at least 1 hour prior to oral specimen collection. Saliva aliquots (1 mL) were stored at –80 °C until RNA and DNA extraction. Total RNA from saliva samples was extracted using the Maxwell® RSC miRNA Tissue kit (Promega) following the manufacturer’s instructions. Initially, 400 µL of saliva samples were added to chilled 200 μL of 1-Thioglycerol per 1 mL of Homogenization Solution, following the instructions of the manufacturer. At the end of the automated extraction, RNA samples were eluted in 60 µL of water. RNA concentration was measured with a Qubit fluorometer using Qubit microRNA assay (Thermofisher). DNA was extracted from 500 µL of saliva with the Maxwell®RSC Stabilized Saliva DNA kit (Promega) according to the manufacturer’s instructions. At the end of the automated extraction, DNA was eluted in 60 µL of water. The DNA quantification was performed with a Qubit fluorometer (Qubit DNA HS Assay Kit; Invitrogen).

### Library preparation for small RNA sequencing

Small RNA sequencing (small RNA-seq) libraries were prepared from RNA extracted from saliva following a protocol for library prep described in^36,37^. Briefly, the NEBNext Multiplex Small RNA Library Prep for Illumina kit (New England Biolabs) was used to convert small RNA transcripts into barcoded complementary DNA (cDNA) libraries. For each library, 100 ng of RNA was processed as starting material. Each library was prepared with a unique indexed primer. Multiplex adapter ligations, RT primer hybridization, RT reaction, and PCR amplification were performed according to the manufacturer’s protocol. After PCR amplification, the cDNA constructs were purified using the Monarch PCR & DNA cleanup Kit (New England Biolabs), following the modifications suggested in the NEBNext Multiplex Small RNA Library Prep for Illumina protocol. Final libraries were loaded on the Bioanalyzer 2100 (Agilent Technologies) using the DNA High Sensitivity Kit (Agilent Technologies) according to the manufacturer’s protocol. Libraries were pooled together (in 35-plex or 40-plex) and further purified with a gel size selection. A final Bioanalyzer 2100 run using the High Sensitivity DNA Kit (Agilent Technologies) was performed to assess DNA library quality with respect to size, purity, and concentration.

The obtained libraries were subjected to the Illumina sequencing pipeline on an Illumina NextSeq500 sequencer (Illumina). Raw and processed sequencing data were deposited on Gene Expression Omnibus (GEO) with the identifier GSE285846.

### Small RNA-Seq data analysis

Small RNA-seq analyses were performed using a Docker-based pipeline to ensure computational reproducibility^38^. Specifically, trimmed reads were mapped against a curated reference of human miRNAs based on miRBase v22.1. BWA algorithm v0.7.12.12 was used for read alignments on miRNA hairpin sequences. Mature miRNA levels were quantified as previously described^38^. In the case of mature miRNAs with identical sequences, the associated read counts were summed. Differential expression analysis was performed using DESeq2 (v1.40.2) with the likelihood ratio test, adjusting for age and sequencing pool. A miRNA was considered differentially expressed (DE) if associated with an adjusted p <0.05 and a median number of reads >10 in at least one study group.

Functional enrichment analysis was performed using RBiomirGS v0.2.19 with default settings and the IWLS method for the rbiomirgs_logistic function. Only validated miRNA-target interactions from miRTarBase 7.0 and miRecord were considered. The enrichment analysis was performed using the gene set libraries (2.cp.reactome.v2023.2, c5.go.bp.v2023.2, c2.cgp.v2023.2) from MSigDB (v2023.2.Hs). A term was considered enriched if it was associated with an adjusted p < 0.05 and at least two target genes. The input for the analysis was the average log2 fold change and the Fisher-combined adjusted P value from the differential expression analyses.

### Library preparation for shotgun metagenomic sequencing

Sequencing libraries were prepared starting from 24 ng of DNA using the Illumina® DNA Prep (M) Tagmentation kit (Illumina), following the manufacturer’s guidelines and as described in^39^. The library pool was subjected to a cleaning step with 0.7x Agencourt AMPure XP beads as described in^40^. Samples were sequenced on a NovaSeq 6000 flow cell (Illumina) at the Italian Institute for Genomic Medicine (IIGM) sequencing facility. Sequencing reads were deposited on SRA with the accession PRJNA1327569.

### Shotgun metagenomics data analysis

The read preprocessing (adapter trimming and removal of low-quality reads), the alignment on PhiX control, and the human genome (hg38) were performed as described in the study by^39,40^, using the pipeline available at https://github.com/SegataLab/preprocessing. The preprocessing steps include: for: i) removal of low-quality reads (quality Q<20), too short fragments (length <75 bp), and reads with two or more ambiguous nucleotides; ii) host and contaminant DNA removal using Bowtie 2.77 (--sensitive-local) for filtering the phiX174 Illumina spike-in and human-aligned reads (hg38 assembly); iii) creation of paired forward and reverse and unpaired reads output files.

Taxonomic profiling was performed with MetaPhlAn 4.1 with the “--statq 0.1” and the ChocoPhlAn vJun23 database^41^. Microbial pathway abundances were estimated using HUMAnN 3.9^42^, applied with default settings and using the full UniRef90 as the reference database. Low-abundant species were removed using the *nearZeroVar* function of the caret R package. Diversity metrics (richness, Inverse Simpson index, Shannon index, and evenness) were computed using the *vegan* R package v2.5-6.1. Differential abundance analysis was performed with SIAMCAT v2.12^43^. The analysis was performed on all taxonomic levels, including the Species-level Genomic Bins (SGBs) profiled using MetaPhlAn 4.1, by removing low-abundant taxa (*nearZeroVar* function of caret R package).

### Statistical and computational analyses

All statistical analyses and graphical representations were performed using R (version 4.5). Statistical analysis between continuous variables was performed using the Wilcoxon rank sum test or Kruskal-Wallis test. Statistical analysis between categorical variables was performed using the chi-square test. Correlation analyses were performed using Spearman’s rank correlation.

Significant microbial pathways were selected using the Wilcoxon rank-sum test, based on the global pathway score. To remove the effect of confounding, a linear model was fitted, adding the age information as a covariate. Only tests with a Benjamini-Hochberg-adjusted p-value <0.1 and retaining the significance (p<0.05) after age correction were considered. Then, individual microbial contributions (genus-level) were calculated using the same method, considering tests with p<0.05.

DIABLO module of MixOmics v.6.32^44^ was used to integrate identified relevant miRNAs and HUMAnN microbial pathways. The analysis was conducted using the projection to latent structures (PLS) method, which applies sparse multiblock partial least squares discriminant analysis for simultaneous integration and variable selection. For the analysis, the first two components were considered, and only significant correlations were selected.

## Results

### Subject characteristics

Police officers were divided into three groups based on SR scores (low-SR, intermediate-SR, and high-SR), and their relative sociodemographic and occupational characteristics are shown in **Table 1**. No significant differences in the distribution of these characteristics were observed among the three study groups stratified by SR levels, except for age. Participants in the high-SR group were, in fact, younger than those in the low- and intermediate-stress groups (p=0.04). This difference may be attributable to the early withdrawal from the cohort by subjects with higher SR levels. Age was then considered as a covariate in the subsequent analyses.

**Table 1.** Sociodemographic and occupational variables of the study population stratified for stress response (SR) categories.

| Variables | Low SR<br>N = 31 | Intermediate SR<br>N = 55 | High SR<br>N = 27 | $p^3$ |
| --- | --- | --- | --- | --- |
| Age, years<sup>1</sup> | 45 ($\pm 5$) | 45 ($\pm 5$) | 42 ($\pm 5$) | **0.04** |
| BMI<sup>1</sup> | 26.1 ($\pm 3.9$) | 26.8 ($\pm 3.6$) | 26.6 ($\pm 3.0$) | 0.40 |
| Education<sup>2</sup> |  |  |  | 0.40 |
| University degree | 1 (3.2%) | 2 (3.6%) | 0 (0.0%) |  |
| High school | 20 (64.5%) | 41 (74.6%) | 23 (85.0%) |  |
| Middle school | 10 (32.3%) | 12 (21.8%) | 4 (15.0%) |  |
| Marital status<sup>2</sup> |  |  |  | 0.60 |
| Unmarried | 15 (48.4%) | 23 (41.8%) | 16 (59.3%) |  |
| Married | 15 (48.4%) | 28 (50.9%) | 9 (33.3%) |  |
| Separated | 1 (3.2%) | 4 (7.3%) | 2 (7.4%) |  |
| Children<sup>2</sup> |  |  |  | 0.30 |
| N | 17 (55.0%) | 27 (49.0%) | 18 (67.0%) |  |
| Y | 14 (45.0%) | 28 (51.0%) | 9 (33.0%) |  |

<sup>1</sup>Mean ( ± standard deviation); <sup>2</sup>n (%); <sup>3</sup>Kruskal-Wallis test; Fisher's exact test; Pearson's Chi-squared test; N: no; Y: yes.

| **Variables** | **Low SR**<br>N = 31 | **Intermediate SR**<br>N = 55 | **High SR**<br>N = 27 | **p<sup>3</sup>** |
| --- | --- | --- | --- | --- |
| Age, years <sup>1</sup> | 45 (±5) | 45 (±5) | 42 (±5) | **0.04** |
| BMI<sup>1</sup> | 26.1 (±3.9) | 26.8 (±3.6) | 26.6 (±3.0) | 0.40 |
| Education<sup>2</sup> |  |  |  | 0.40 |
| University degree | 1 (3.2%) | 2 (3.6%) | 0 (0.0%) |  |
| High school | 20 (64.5%) | 41 (74.6%) | 23 (85.0%) |  |
| Middle school | 10 (32.3%) | 12 (21.8%) | 4 (15.0%) |  |
| Marital status<sup>2</sup> |  |  |  | 0.60 |
| Unmarried | 15 (48.4%) | 23 (41.8%) | 16 (59.3%) |  |
| Married | 15 (48.4%) | 28 (50.9%) | 9 (33.3%) |  |
| Separated | 1 (3.2%) | 4 (7.3%) | 2 (7.4%) |  |
| Children<sup>2</sup> |  |  |  | 0.30 |
| N | 17 (55.0%) | 27 (49.0%) | 18 (67.0%) |  |
| Y | 14 (45.0%) | 28 (51.0%) | 9 (33.0%) |  |

<sup>1</sup>Mean ( ± standard deviation); <sup>2</sup>n (%); <sup>3</sup>Kruskal-Wallis test; Fisher's exact test; Pearson's Chi-squared test; N: no; Y: yes.

### Salivary miRNAs are differentially expressed in different stress response groups

miRNome profiled in saliva samples by small RNA-seq generated an average of 44,243 reads aligned on human miRNA annotations and an average of 254 miRNAs detected in each sample (range:186-549) (**Supplementary Table 1A**). The most abundant salivary miRNA identified was miR-148a-3p, followed by miR-526a-5p, miR-520c-5p, miR-518d-5p, miR-3135a-3p, miR-223-3p, and miR-1246. An age-adjusted differential expression analysis was performed by comparing high-vs. low-SR, as well as intermediate- vs. low- and intermediate- vs. high-SR groups. In the comparison high- vs. low-SR, 18 miRNAs displayed significant different levels among groups, with four (miR-10400-5p, miR-1290, miR-6074-5p, and miR-9902) and fourteen (miR-203a-3p, miR-7-5p, miR-143-3p, miR-181a-5p, miR-142-5p, miR-223-5p, let-7f-5p, let-7g-5p, miR-26b-5p, miR-27a-3p, miR-30e-5p, miR-26a-5p, miR-142-3p, and miR-21-5p) miRNAs characterized by higher and lower levels, respectively, in the high-SR group (**Figure 1A** and **Supplementary Table 1B**).

**Figure 1.**
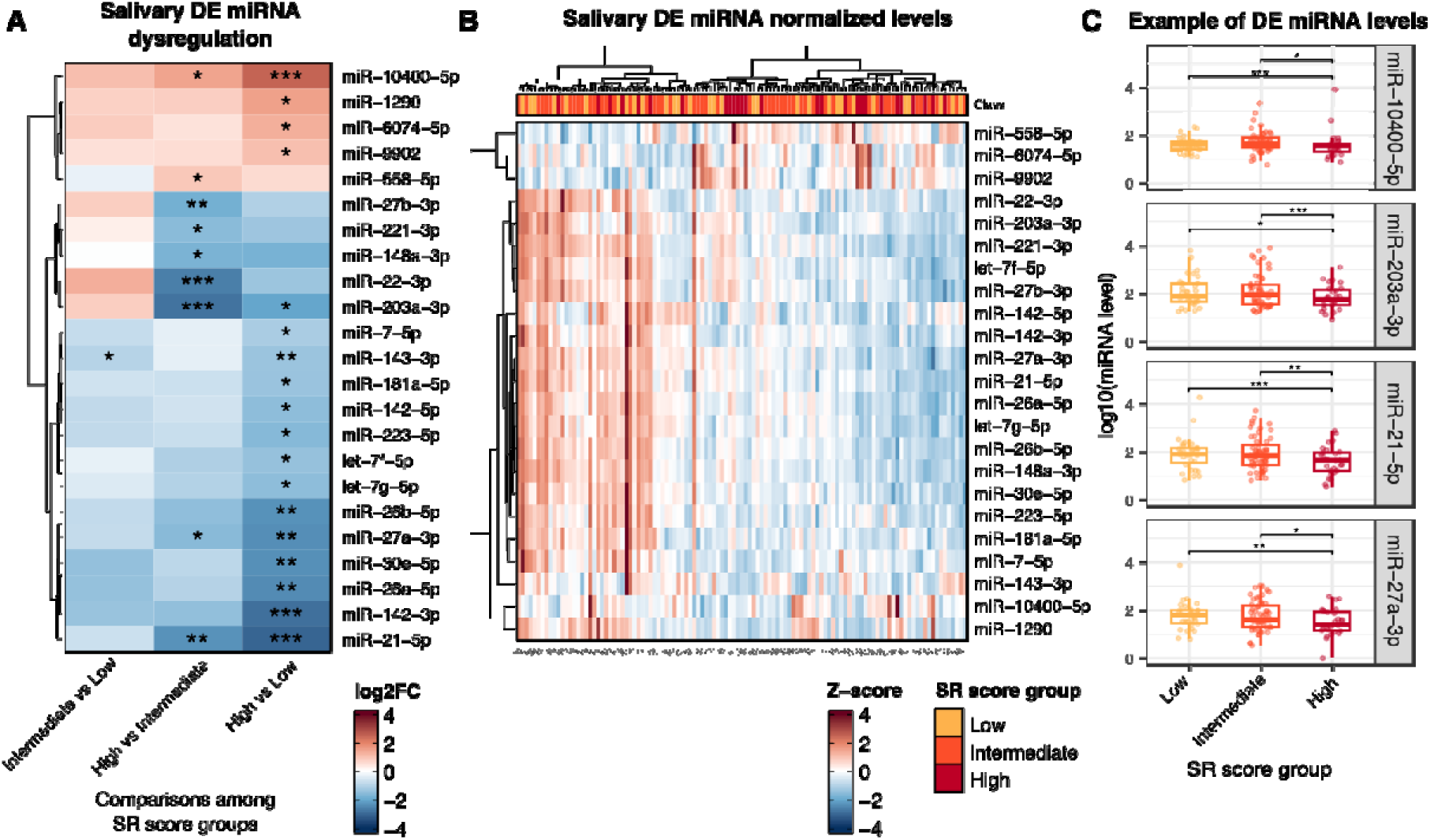
Analysis of salivary DE miRNA profiles. Heatmap of the log2FC (**A**) and Z-score (**B**) of the DE miRNAs observed in at least one comparison. In (**B**), for each participant, the SR (Stress Response) score level is reported. Adjusted p-value from DESeq2 analysis: ^∗∗∗^adj. p < 0.001; ^∗∗^adj. p < 0.01; ^∗^adj. p < 0.05. **C**) Boxplot reporting the levels of DE miRNAs with altered expression in both intermediate and high SR subjects. ***adj. p < 0.001; **adj. p < 0.01, *adj. p < 0.05.

On the other hand, only miR-143-3p levels significantly decreased in the intermediate group when compared with the low-SR group, while nine miRNAs (miR-10400-5p, miR-558-5p, miR-27b-3p, miR-221-3p, miR-148a-3p, miR-22-3p, miR-203a-3p, miR-27a-3p, miR-21-5p) were associated with significantly different levels between the intermediate and the high-SR group. In this last set, the levels of miR-10400-5p and miR-558-5p were increased in the high-SR group, and those of miR-27b-3p, miR-221-3p, miR-148a-3p, miR-22-3p, miR-203a-3p, miR-27a-3p, and miR-21-5p decreased (**Figure 1A-B**). Interestingly, the levels of four miRNAs (miR-10400-5p, miR-203a-3p, miR-27a-3p, and miR-21-5p) were commonly altered across multiple comparisons and showed a progressive increase or decrease from the low- to the high-SR group (**Figure 1C**).

### miRNA target enrichment analysis

Functional analysis of the identified DE miRNA target genes (**Figure 2A** and **Supplementary Table 1C-D**) showed an enrichment in genes involved in apoptosis-related processes, cellular response to stress, negative regulation of gene expression, responses to abiotic stimuli, positive regulation of catabolic processes, and macromolecule catabolic processes. Network analysis of the miRNA-target interactions supported by the highest number of evidence showed miR-21-5p as the main miRNA hub, followed by miR-223-5p, and miR-27b-3p/miR-27a-3p. Several genes were targeted by miRNAs with decreasing levels in the high-SR group, including *PTEN*, *PITHD1*, *MDM2*, *MAT2A*, *KPNA2*, *FIGN*, *CDKN1B*, *CCND1*, and *BCL2* (**Figure 2B** and **Supplementary Table 1E**).

**Figure 2.**
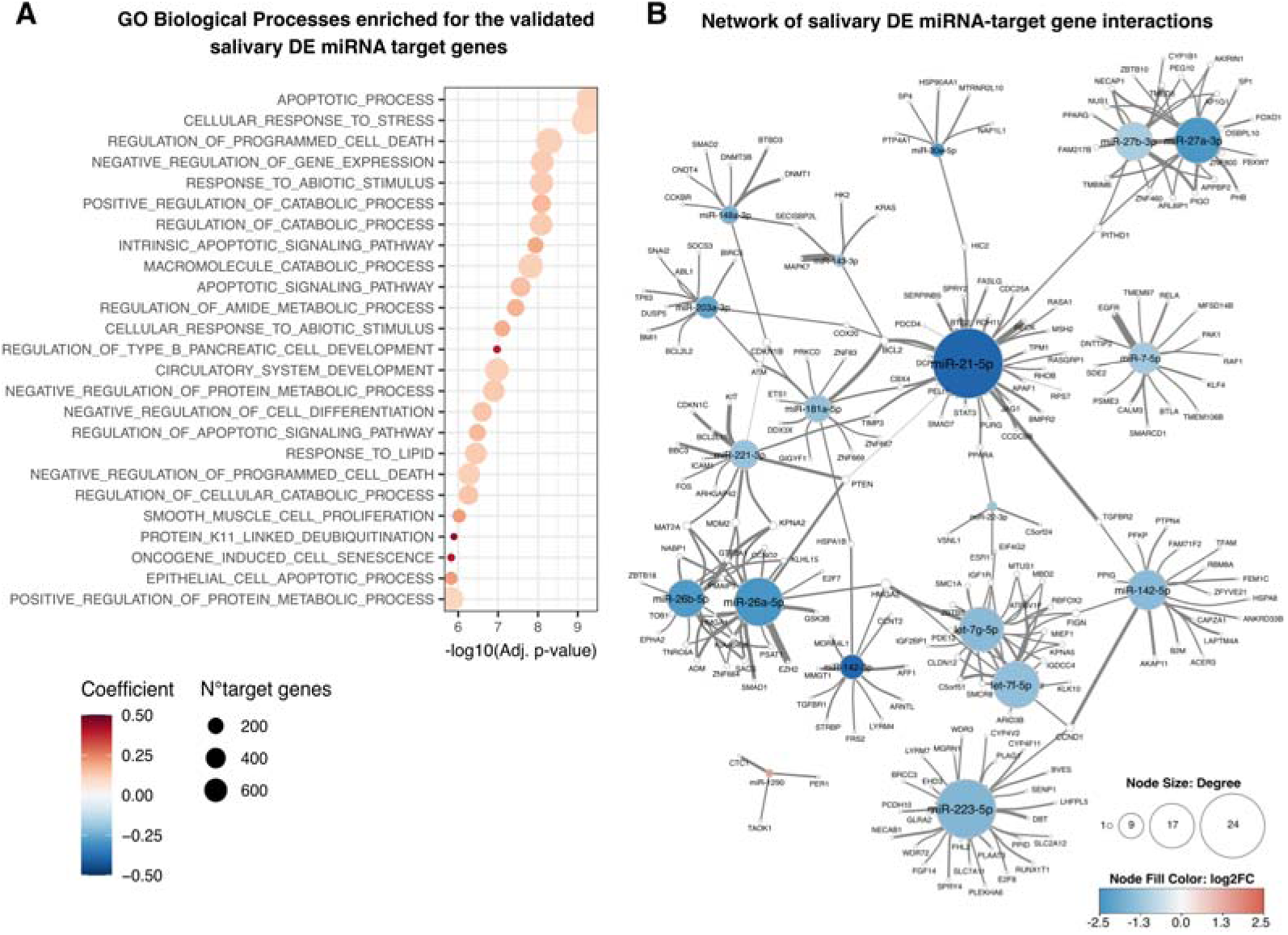
Functional target enrichment analysis of DE salivary miRNAs. **A**) Functional enrichment analysis results from RBiomirGS analysis. The dot size is proportional to the number of target genes, while the color code represents the pathway activation coefficient. Red coefficients indicate pathways predicted to be activated by targeting miRNA expression changes. **B**) Network representation of the miRNA-target interaction supported by at least four independent publications. The node size is proportional to the total node degree, while the color code reflects the expression change (log2FC) computed between the low- and high-SR groups. Edge tightness is proportional to the amount of supporting evidence.

### Salivary metagenome profiles

Shotgun metagenomic sequencing was performed on the same set of samples analyzed for miRNAs. After removal of host DNA content, an average of 18.3±8.4 million reads were obtained, of which an average of 26.53% were assigned to microbial annotations, resulting in 237.0±82.9 SGBs detected across samples (**Supplementary Table 2A**). Microbial diversity analysis among the three study groups shows no significant differences in alpha diversity (Richness, Shannon index, Inverse Simpson index and Evenness) (**Figure 3A** and **Supplementary Figure 1A**), nor in the beta diversity (PERMANOVA p>0.05) (**Figure 3B**). Since functional microbial alterations cannot be captured solely at taxonomic levels, a taxon-set enrichment analysis was performed to explore candidate functional alterations in microbial communities. The analysis showed significant enrichments (p<0.05) in dimethylamine-producing (p<0.01) and fructose- and gelatinase-utilizing taxa (**Figure S1B**). Across taxonomic levels, the *Lachnospiraceae unclassified* genus abundance decreased in the high SR group (adj. p<0.05) (**Figure 3C** and **Supplementary Table 2B**). Despite not being significant after multiple-testing correction, some SGBs showed significant differential abundance in the high-SR groups (**Figure 3D**, **Figure S1C** and **Supplementary Table 2B**).

**Figure 3.**
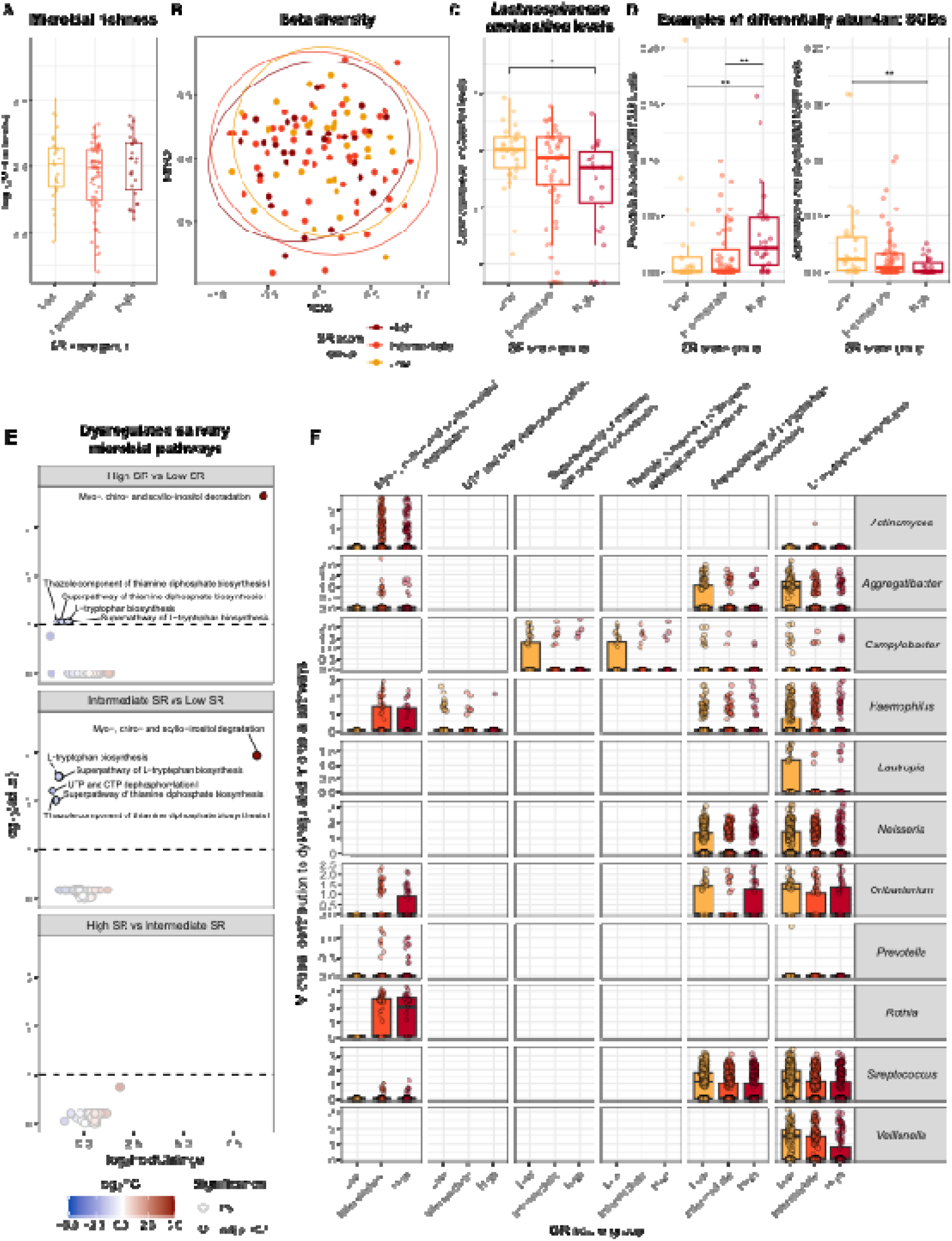
Analysis of salivary microbial communities. **A**) Box plots reporting the microbial richness across SR groups. **B**) Nonmetric multidimensional scaling plot based on inter-subject Bray-Curtis dissimilarity. **C**) Box plots reporting the *Lachnospiraceae unclassified* abundance across SR groups. **D**) Box plots reporting the levels of differentially abundant oral SGBs across SR groups. The box and color reflect the SR group. **p<0.01, *p<0.05 by Wilcoxon rank-sum test. **E**) Volcano plots of the microbial pathways differential analysis. On the X-axis, the log2FC is reported, while on the Y-axis, the significance (adj. p-value) is shown. The black dashed line indicates the adj. p-value threshold of 0.1, and the black dot border highlights the significant ones. **F**) Boxplots reporting the contribution of different microbial genera to each identified pathway (contribution score, Y-axis).

The analysis of the microbial pathway abundance also identified six pathways differentially enriched among the study groups, whose levels were not affected by age (**Figure 3E** and **Supplementary Table 2C**). Among them, the *Myo-, chiro- and scyllo-inositol degradation* (PWY-7237) was characterized by the most significant increase in the high-SR group, while *L-tryptophan biosynthesis* (TRPSYN-PWY) was characterized by the most significant decrease. Conversely, no pathways were significantly altered between the high- and intermediate-SR groups (**Figure 3E** and **Supplementary Table 2C**). Investigation of microbial species contributing to the identified pathways showed that the increase of the *Myo-, chiro- and scyllo-inositol degradation* (PWY-7237) was mainly driven by Actinomyces, Haemophilus, and Oribacterium bacteria (**Figure 3F** and **Supplementary Table 2D**). On the other hand, in the high-SR group, the decrease in genes involved in *L-tryptophan biosynthesis* was primarily driven by a reduction in Aggregatibacter, Neisseria, Streptococcus, and Veillonella.

### Integrative analysis of miRNA and metagenomic profiles

To identify the subset of molecular features (salivary miRNAs, SGBs, and microbial pathways) with the highest discriminatory potential for the study groups, an integrative analysis using DIABLO was performed. The analysis showed that the microbial pathways were associated with the greatest potential for separating the high- from the low-SR groups (**Figure 4A**). Among the most informative features from each molecular layer, miR-7-5p, miR-1290 and miR-142-3p were the most represented miRNAs, whereas *Myo-, chiro-, and scyllo-inositol degradation* (PWY-7237), *UTP and CTP dephosphorylation I*, and the *Thiazole component of thiamine diphosphate biosynthesis* (PWY-6892) were the most informative microbial pathways (**Figure 4B**). The impact of these microbial pathways on separating the study groups was clearly highlighted by the multi-omic clustering analysis, which showed that the high- and intermediate-SR groups were distinct from the low-SR group (**Figure 4C**).

**Figure 4.**
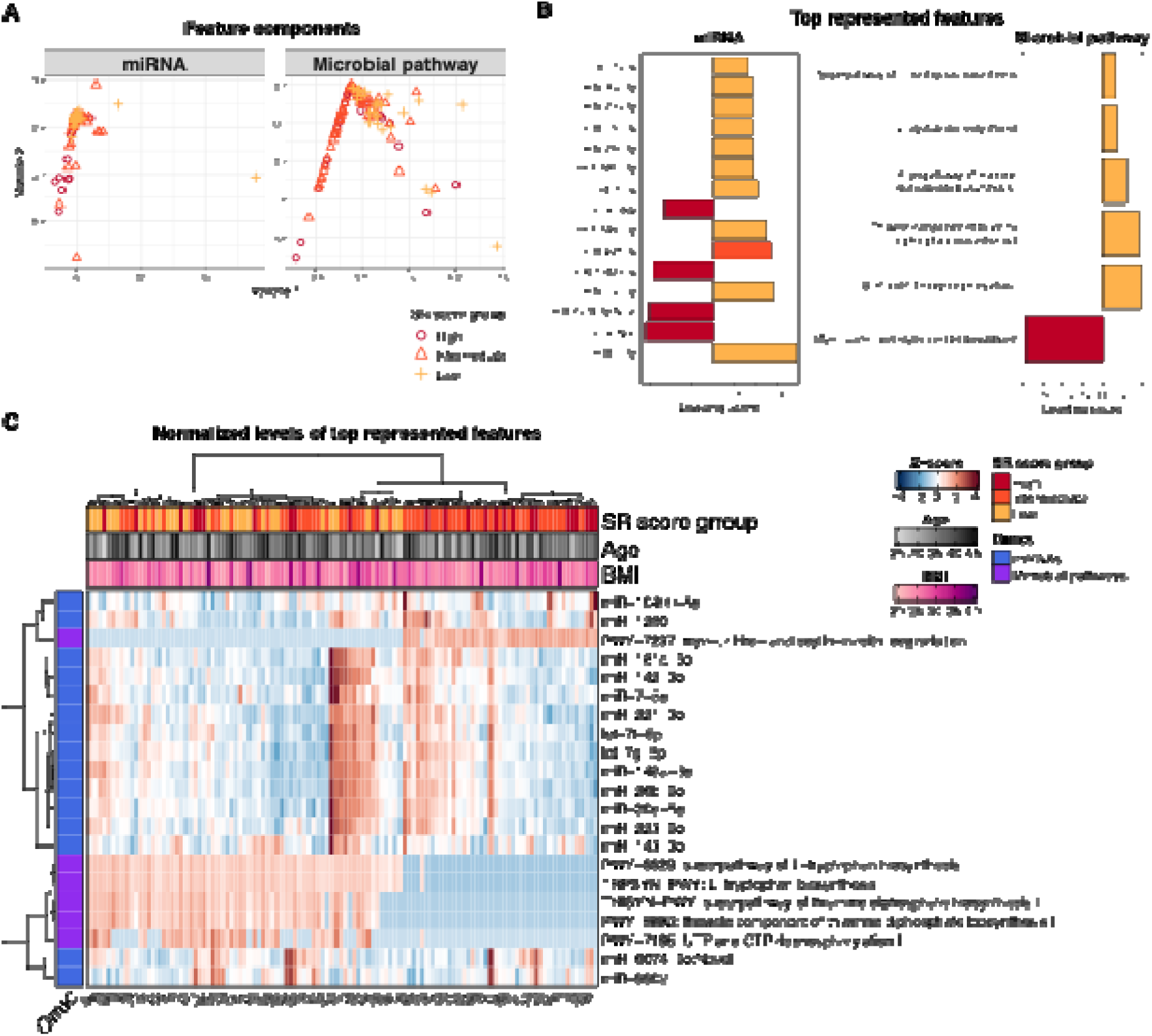
Integrative analysis of salivary miRNA and microbial profiles. **A**) PLS plot reporting Variate 1 and Variate 2, which represent the components capturing the highest covariance between omics features and SR groups. **B**) Bar plots reporting the most informative features in each omics dataset. The X-axis reports the contribution of each feature (loading score), while the color reflects the SR group. **C**) Heatmap of the unsupervised clustering of Z-score-transformed levels of the top correlated features. The single omics are highlighted with a specific color, while a different color key highlights the subject’s SR group. For each subject, the age and BMI value are also reported.

## Discussion

In the present study, salivary miRNA and microbial profiles were investigated in a cohort of otherwise healthy police officers, well characterized for their occupational exposure and stress responses. Despite the small sample size, the data confirmed that psychosocial stress is associated with different salivary miRNA and microbial profiles. For the first time, it was shown that high levels of psychosocial SR are characterized by a specific salivary miRNA expression pattern and distinct microbial pathways that show dose-dependent increases or decreases compared with those of resilient subjects with low SR. We chose saliva as a sampling matrix because it is not only non-invasive and easily collected and managed, but it has also proven useful for assessing miRNAs and microbial species as biological markers across different pathophysiological conditions, including stress^16,45^.

miRNAs have emerged as an integral part of SR for many years^6,8,10,46,47^, and their role as biomarkers of both acute^9,11,12,45^ and chronic^9,11^ psychological stress has been reported, either in experimental or naturalistic settings. Although there is growing interest in the relationship between miRNAs and stress, occupational studies in this field are few. Literature demonstrated that work-related stress is associated with an increased risk of mental and physical issues^48^. To the best of our knowledge, there are no studies of miRNA profiling conducted on police officer cohorts in which potential confounding variables were largely controlled, and SR was monitored for several years, thereby demonstrating longitudinal stability of individual differences in physiological responses to work-related stressors.

The observed alterations in miRNAs between high- and low-SR groups either confirmed known stress-associated miRNA differences or identified a new set of miRNAs associated with distress. Notably, miR-21-5p expression has already been linked to human psychological stress, and, consistent with our results, salivary miR-21-5p levels decreased under an acute stress test (the Trier social stress test)^45^, and in whole blood under a stressful event^49^. miR-21-5p is a well-known regulator of stress-related response, apoptosis and inflammation, and it is frequently overexpressed in neoplastic, inflammatory, cardiovascular, and neurodegenerative diseases^50^. Although the mechanisms leading to miR-21-5p decrease in highly distressed subjects are presently unknown, we can speculate a role for sustained sympathetic activation with miR-21-5p downregulation. Indeed, the stress mediator adrenaline, released upon sympathetic nervous system activation, reduces miR-21 expression in human cells^51^. An association has also been found between salivary alpha-amylase, a marker for sympathetic activation, and salivary miR-21 levels under acute psychological stres^45^.

Salivary miR-26a-5p and miR-26b-5p levels were also decreased in high-SR police officers which have been previously associated with psychological stress^9,45^. miR-26a-5p is highly expressed in the central nervous system, where it regulates synaptic plasticity, neuronal morphogenesis, neurotransmission, and axon regeneration^52^. Studies have reported increased circulating and salivary levels of miR-26a-5p and miR-26b-5p under stress conditions^9,45^. However, there may be obvious differences in types, duration, and intensity of stressors between different studies, and in the case of the present investigation, any observed differences in miRNA levels and patterns were associated with (and potentially driven by) long-standing high stress, which may imply unique molecular adaptations reflecting established resilience or vulnerability phenotypes.

A decrease in salivary let-7g-5p and let-7f-5p levels was also observed in the high-SR group. These miRNAs are members of the let-7 family and have previously been reported to decrease in whole blood in neurodegenerative diseases^53^. In contrast, let-7g-5p levels increased in subjects with chronic fatigue syndrome, sleep disorders, mood swings, headaches/migraines, depression, anxiety, and other likely stress-related diseases^11^. Another informative miRNA emerging from our analysis was miR-142-3p, which has been previously reported to mediate neuroinflammation-induced synaptic dysfunction in mice^54^, and it is downregulated in immune cells of aged animals with associated elevation of proinflammatory factors^55^. Notably, the expression levels of miR-142-3p-target gene *AFF1* were upregulated in microglia of stressed mice compared with controls, as part of a transcriptional program regulating catabolism and macromolecule biosynthesis, in line with increased metabolism and energy demand under chronic stress^56^.

A previous study reported that miR-27a-3p levels decreased in the blood of rats displaying vulnerability (increased anxiety-like behavior) to chronic stress (social defeat) compared with resilient rat^57^. Other miRNAs with decreasing salivary levels in the high SR group in this study were miR-181a-5p, miR-7-5p, and miR-143-3p; the first two have been previously reported to be involved in various types of cellular stress, including endoplasmic reticulum stress^58^ and oxidative stress^59^. Conversely, miR-143-3p has been involved in the occurrence of depressive-like behaviors in mice, mediating alterations in synapse density in the hippocampus^60^.

In the comparison of the high SR group with the intermediate SR group, a significant specific deregulation was found in five miRNAs: miR-558-5p (with increasing levels), miR-27b-3p, miR-221-3p, miR-148a-3p, and miR-22-3p (all with decreasing levels). Moreover, a few miRNA sets were common to those altered under high versus low SR and followed the same pattern: miR-203-3p, miR-27a-3p, miR-21-5p (decreasing), and miR-10400-5p (increasing). This pattern suggests early perturbation of salivary miRNAs with an intermediate level of work-related SR. Despite, the functions of these miRNAs in SR are unknown, serum miR-221-3p and miR-27a-3p levels were increased in subjects with posttraumatic stress disorder^61^.

In contrast to the evidence on the relation between psychosocial stress and gut microbiota^14^, a few preclinical and clinical studies have found that psychological stress can induce changes in the oral microbiome^16,62^. Our metagenomic analysis showed no significant differences in ecological metrics of salivary microbiota composition among subjects, with only a few SGBs exhibiting differential abundances across the study groups (p<0.01). Specifically, *Prevotella baroniae|SGB1533*, a species that has been previously reported to play a role in stress-related disorders, including anxiety and depression^63^, was more abundant in the high-SR group compared to the low-SR group. Accordingly, altered oral levels of the Prevotella genus were reported in subjects with psychological stress^64^.

Analysis of microbial pathways showed a clear SR group separation based on putative metabolic processes: the *Myo-, chiro- and scyllo-inositol degradation* pathway had an increased activity in high-SR groups, while the *L-tryptophan biosynthesis*, *Superpathway of thiamine diphosphate biosynthesis*, and *Thiazole component of thiamine diphosphate biosynthesis* were decreasing with the SR severity. Inositol is a cyclic sugar alcohol that acts as a precursor of cellular secondary messengers and plasma membrane components, whose metabolism is related to microbial fermentation^65^. Myo-inositol is the most abundant inositol in mammalian cells, and it is a pivotal regulator of neuronal connection formation during development^66^. Animal studies have shown an increase in inositol brain levels in chronic psychosocial stress^67^. However, the microbial degradation of inositol has been mainly characterized for the gut microbiota and its role in energy conservation and lipid synthesis, with both favorable and adverse metabolic outcomes associated with increased inositol metabolism^68^.

In parallel, in the high-SR group, a decrease in microbial genes involved in the biosynthesis of the essential amino acid L-tryptophan was observed. The microbiota can influence L-tryptophan metabolism via several pathways^69^, with consequences for both the host and bacterial physiology. L-tryptophan is the precursor for the biosynthesis of the gastrointestinal and neuroendocrine transmitter serotonin (or 5-hydroxytryptamine). Peripheral serotonin is predominantly synthesized by colonic enterochromaffin cells and exerts multiple physiological functions, including regulation of gut motility and fluid secretion, energy metabolism, cardiac contraction, vascular tone, coagulation, and the immune response^70^. Moreover, gut-derived serotonin communicates bidirectionally with the brain’s serotonergic system via the gut–brain axis^71^. Certain taxa, including *Corynebacterium*^72^, possess the genomic potential to synthesize serotonin, whereas others can regulate host serotonin levels through microbiota-derived metabolites or cell components that affect serotonin biosynthesis in enterochromaffin cells. Accordingly, a reduction in *Corynebacterium* was observed in both the human saliva microbiome and the rat oral microbiome of individuals under high stress^73^.

L-tryptophan can be metabolized by the gut microbiota into the neurotransmitter tryptamine, as well as indole and its derivatives, which are ligands of the aryl hydrocarbon receptor and participate in immune and intestinal homeostasis^69^. Recent findings have disclosed a neuroprotective role for a microbiome-derived indole against chronic psychosocial stress in a mouse model ^74^. Tryptophan can also be degraded by host immune and intestinal cells and by some bacterial species through the kynurenine pathway, thereby generating kynurenine and downstream products, such as kynurenic acid and quinolinic acid, which play roles in inflammation, immune responses, and neurobiological functions^69^. Several reports have shown that chronic stress and depression trigger the kynurenine pathway through the induction of inflammation, thus diverting tryptophan metabolism from the production of serotonin or indoles to the synthesis of neuroactive/neurotoxic metabolites^75,76^. Although chronic stress is associated with dysregulated L-tryptophan metabolism in both the central nervous system and the periphery, with a contributory role for an altered gut microbiome^77^, the implications of changes in oral microbiota composition and metabolism remain unclear^78^.

Thiamine diphosphate is the active coenzyme form of thiamine (vitamin B1), which is crucial for energy metabolism and, consequently, for various host and bacterial metabolic and immunoregulatory processes, as well as for neurotransmitter synthesis and protection against oxidative stress. Certain bacterial strains have long been known to contribute to vitamin B1 synthesis and availability for both the host and bacteria^79^. The observed reduction in the metabolic pathway related to thiamine diphosphate biosynthesis in the oral microbiome of the present study could be of interest in the context of chronic stress, since studies have suggested a (neuro)protective role for thiamine against stress, anxiety, depression and mental health in general^80^. Moreover, a decreased thiamine metabolic pathway can negatively impact the microbiome itself in a vicious cycle^81^, thus potentially contributing to the reshaping of the microbiota as observed in response to stress.

Our findings are unique because they focus on the oral microbiome, a less-explored niche compared with the gut microbiota. Therefore, although our study revealed specific differences in oral microbial composition and function, additional investigations are needed to elucidate the potential roles of the oral-brain axis and intrinsic neural, immune, and microbial pathways in regulating SR. We cannot determine whether the differences in salivary miRNAs and microbial pathways observed here represent a compensatory protective response to stress or reflect maladaptive physiological and psychological responses. It is possible that both conditions play out in work-related SR.

The sample size was among the limitations of this study, although it was comparable to that of other studies on miRNAs and microbiota profiling and SR. However, this limitation was balanced by the advantage of having studied a population exposed to intense levels of occupational stress, with repeated control for possible confounding factors (diet, physical activity). Monitoring the SR - and the associated psychological and clinical outcomes - for many years has allowed the measurement of the chronic occupational SR, data that has so far been absent in the literature. Another limitation of this study was the lack of women in the cohort. We cannot, therefore, determine the influence of the hormonal environment on miRNAs and microbial species, nor can we study their relationships with stress hormones or inflammatory factors. Moreover, our study cohort was not screened for overall oral health and dietary habits, which introduces a further limitation related to major confounders. Larger prospective studies that include female workers would allow researchers to assess the clinical value of the observed changes induced by psychosocial stress.

On the other hand, the strengths of the study derived from 1) the use of a high-throughput approach (small RNA-seq and metagenomics) to explore salivary miRNA and microbial profiles, which is not based on a priori selection of candidate features and allows the identification of novel miRNAs/taxa associated with response to stress; 2) the use of an homogeneous study sample, with all recruited policemen engaged in maintaining law and order, and exhibiting, at recruitment, levels of stress comparable to the baseline ones (several years before), thus representing a naturalistic setting for studying true chronic stress resilience or vulnerability.

The finding that an unfavorable response to work-related psychosocial stress may be associated with specific miRNA and microbial signatures is novel and offers a new perspective on the pathogenic mechanisms of stress and their associated risks and outcomes. These alterations may reflect epigenetic differences and dysbiosis associated with work-related stress and may be accompanied by related physiological and psychological perturbations. The study also provides evidence that miRNAs from human saliva and oral microbiome pathways could offer insights into functional host-microbiome crosstalk, whose alterations may be involved in neuropsychiatric disorders.

## Declarations

### Ethics approval and consent to participate

The study protocol was approved by the Ethics Committee of the Università Cattolica del Sacro Cuore of Rome, Italy (approval n. 285, 16 July 2020), and was conducted in accordance with the ethical standards of the Declaration of Helsinki. All participants provided their written informed consent to participate in the study.

### Consent for publication

Not applicable.

### Availability of data and materials

The datasets generated and analyzed in the current study are available in GEO and SRA under the accession numbers GSE285846 (small RNA-Seq) and PRJNA1327569 (shotgun metagenomics).

### Authors’ contributions

SG: conceptualization, methodology, project administration, supervision; NM: conceptualization, methodology, supervision; BP and ST: methodology, investigation; GF and AC: formal analysis, data curation, visualization; FC: supervision; EG: investigation, writing - original draft; AN: resources, funding acquisition, supervision. All authors read and approved the final manuscript.

## Acknowledgements

This study was made possible thanks to the cooperation of the Italian Police Force (Ministry of the Interior), which supported the data collection. In particular, thanks to Dr. Francesca Cozzone and all the police officers assigned to the “VI Reparto Mobile” of Genoa, who participated in the study.

## Supplementary information

**Supplementary Table 1**

**A)** Alignment statistics from small RNA-seq data for all samples analyzed in this study. **B)** Differential expression analysis of the small RNA-Seq data. **C)** List of the validated genes targeted by miRNAs that were differentially expressed in this study. **D)** miRNA-target gene interactions supported by literature. **E)** Pathway enrichment analyses of differentially expressed miRNAs as derived from RBiomirGS v0.2.19.

**Supplementary Table 2**

**A)** Alignment statistics and information of the saliva shotgun metagenomic sequencing data. **B)** Differential abundances of microbial taxa between the stress response groups. **C)** List of the microbial pathways that were significantly altered between the stress response groups. **D)** List of pathways and relative genus contribution.

**Supplementary Figure 1**

**Figure S1.** Analysis of salivary metagenomic profiles. **A)** Box plots reporting alpha-diversity metrics across SR groups. The significance was evaluated by the Wilcoxon Rank-Sum test. **B**) Bar plot reporting the taxon-specific enrichment analysis (TSEA) results. The X-axis reports the terms, while the Y-axis reports the significance of the analysis. The bar color reports the set class. **C)** Heatmap reporting the log2FC (left) and Z-score (right) of all the differentially abundant SGBs in at least one comparison. For each subject, the SR group is reported. *p < 0.05; **p < 0.01.

