## Supplementary material for "Salivary miRNA and microbial profiles reflect different responses to psychosocial stress": Figure S1

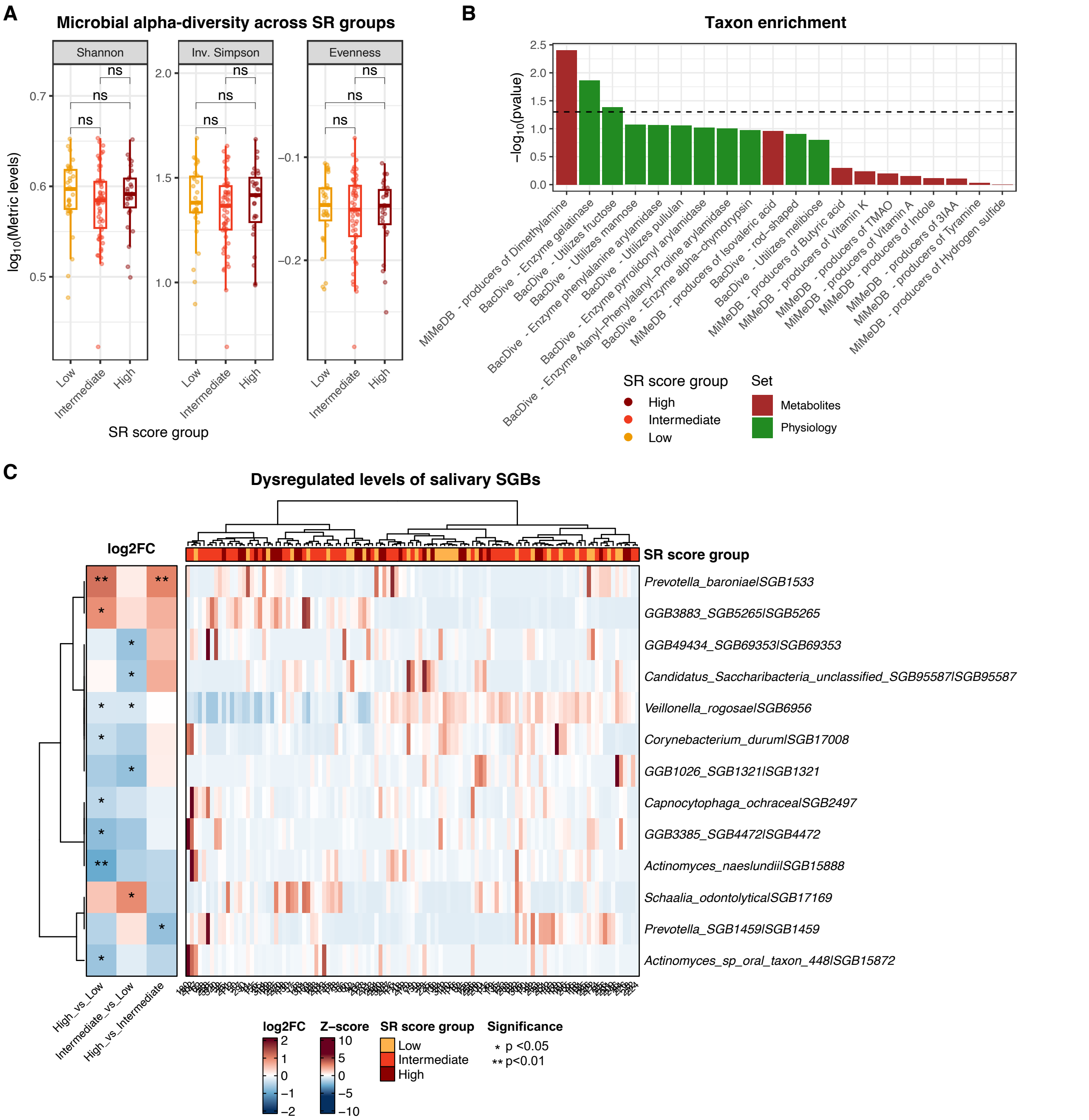


**Figure S1.** Analysis of salivary metagenomic profiles. **A)** Box plots reporting alpha-diversity metrics across SR groups. The significance was evaluated by Wilcoxon Rank-Sum test. **B)** Bar plot reporting the taxon-specific enrichment analysis (TSEA) results. The X-axis reports the terms, while the Y-axis reports the significance of the analysis. The bar color reports the set class. **C)** Heatmap reporting the log2FC (left) and Z-score (right) of all the differentially abundant SGBs in at least one comparison. For each subject, the SR group is reported. $^*p < 0.05$; $^{**}p < 0.01$.
